# A domestication change at *PvMYB26* in common bean sheds light on the origins of Middle American agriculture

**DOI:** 10.1101/2025.05.15.654381

**Authors:** Burcu Celebioglu, Jayanta Roy, Andrew Farmer, Stephanie English, Xingyao Yu, Xiaosa Xu, Phillip E. McClean, Paul Gepts, Travis Parker

## Abstract

Domestication imposed radically different selection pressures on plants, eventually transforming them into the cultivated forms that support global populations today. Here, we investigate the loss of seed dispersal via pod shattering during common bean (*Phaseolus vulgaris* L.) domestication. We identified an eight-kb deletion eliminating the transcription start site and promoter of the candidate gene *PvMYB26*. Mutants express *PvMYB26* at < 1% of the level of wild types and produce 44% less pod lignin. Whole-genome sequencing revealed that the mutation is nearly diagnostic for domestication status among Middle American common bean, indicating its importance in domestication. We also identified a high-frequency *PvMYB26* frameshift/premature stop mutation unique to Andean domesticates. Wild haplotypes most like Middle American domesticates are found in eastern Jalisco, Mexico. Our results suggest that West-Central Mexico was the site of common bean domestication and suggest that this region may have been important in the rise of Middle American agriculture.

## Introduction

Plant domestication transformed the structure and adaptation of domesticated species, leading to a revolution in human evolution and social organization^1-3^. Domestication imposed novel selection pressures on plants, whereby mutations that would be deleterious in the wild became advantageous in the new cultivated environments. For example, elaborate seed dispersal mechanisms are essential to the survival of wild plants, but these became disadvantageous among seed-propagated species in cultivated agroecosystems. Consequently, there was strong selection against seed dispersal during plant domestication, across diverse taxa such as cereals^4^, grain legumes^5^, *Brassicas*^6^, and other seed crops^7^.

Seed dispersal in many legumes (Fabaceae) is mediated by pod shattering, with dry pods violently dehiscing at maturity due to pod walls rapidly coiling under significant torsional forces^5^. This trait was problematic for harvest and yield in cultivated environments, so reduced pod shattering became a core domestication syndrome trait in the family^8-10^, along with loss of seed dormancy^10-13^, and loss of pod shattering occurred up to 40 times in parallel domestications across the family^5^. Among these domesticated species, common bean (*Phaseolus vulgaris* L.) is the most important for direct human nutrition^14-16^. *P. vulgaris* was independently domesticated in Middle America and the Andes from gene pools that had diverged more than 100,000 years ago^17-20^, before human habitation of the Americas. In the Andean gene pool, variation in pod dehiscence has been fine-mapped to *PvMYB26*, previously postulated to be an improvement trait in snap beans^7,21,22^. To date, no causal mutations have been described in this gene in common bean, and the basis of reduced pod shattering during the initial domestication of the species is unknown.

Similarly, unresolved questions remain regarding the origins of agriculture in Middle America. These agricultural systems were founded around the staple crops maize (*Zea mays*), beans (*Phaseolus* spp.), and squash (*Cucurbita* spp.), known collectively as “the three sisters”. Within common bean, contrasting hypotheses regarding the geographic domestication center have been proposed: 1) the West-Central Mexico hypothesis, particularly in the state of Jalisco^23,24^, and more recently, 2) the Southern Mexico hypothesis, in the state of Oaxaca^25^. Since *Phaseolus* beans were particularly important during agricultural intensification and are an integral component of the Middle American crop assemblage, assessing their domestication origin is valuable for understanding the rise of agriculture in Middle America. Domestication alleles are rarely exchanged between wild and domesticated gene pools^26-28^, thereby offering a unique opportunity to determine crop origins without the effects of genetic exchange that occurs broadly in the rest of the genome^26,29-31^.

Here, we investigate the domestication of this critical New World staple crop through genetic mapping of two populations, whole-genome sequencing analysis of 323 novel wild and domesticated accessions, characterization of gene expression through qPCR and RNA-seq, developmental and biochemical screens, and geographic information system analysis. Our results shed light on the genomic basis of a vital domestication trait and the origins of agriculture in Middle America.

## Results

### Variation in pod traits maps to *PvMYB26* and *PvPDH1*

Both measured pod traits (number of pod valve twists and proportion of pods shattered) were significantly correlated in both the UC Canario 707 x Wild PI 638850 (CxW) and Orca x Wild PI 638850 (OxW) populations (*P* < 2*10^-16^). Measurement of pod twists yielded more significant maximum LOD scores than direct measurement of pod shattering (Fig. 1, Supplementary Table 1). The most significant QTL for pod twists was mapped to Pv05 at 38,633,111 bp (LOD = 33.2) in the OxW population and 38,800,513 bp (LOD = 10.6) in the CxW population. The candidate gene *PvMYB26* (chromosome Pv05: 38.6 Mb; G19833 v2.1 gene model *Phvul.005G157600*) is located between the flanking markers of the detected QTL in both populations (Fig. 1b,e, Supplementary Table 1). A secondary locus for pod twists was identified on Pv04 in the OxW population. For pod shattering, two significant QTL were detected in the OxW population (Pv03: 49,199,252 bp, LOD=15.9; Pv05: 38,567,439 bp, LOD=10.4), with the candidate genes *PvPDH1* and *PvMYB26* falling between the peak markers of each QTL, respectively. The CxW population had three significant shattering QTL, mapping near *PvPDH1*, *PvMYB26*, and an additional locus on Pv03 (Supplementary Table 1). In both traits and both populations, no significant interactions were identified between loci, suggesting additive rather than epistatic relationships between them (fitqtl interaction *P* > 0.05). RILs inheriting the wild-type allele had increased trait values in all cases. Each significant QTL explained 3.5-39.8% of phenotypic variance (Supplementary Table 1).

**Fig. 1.**
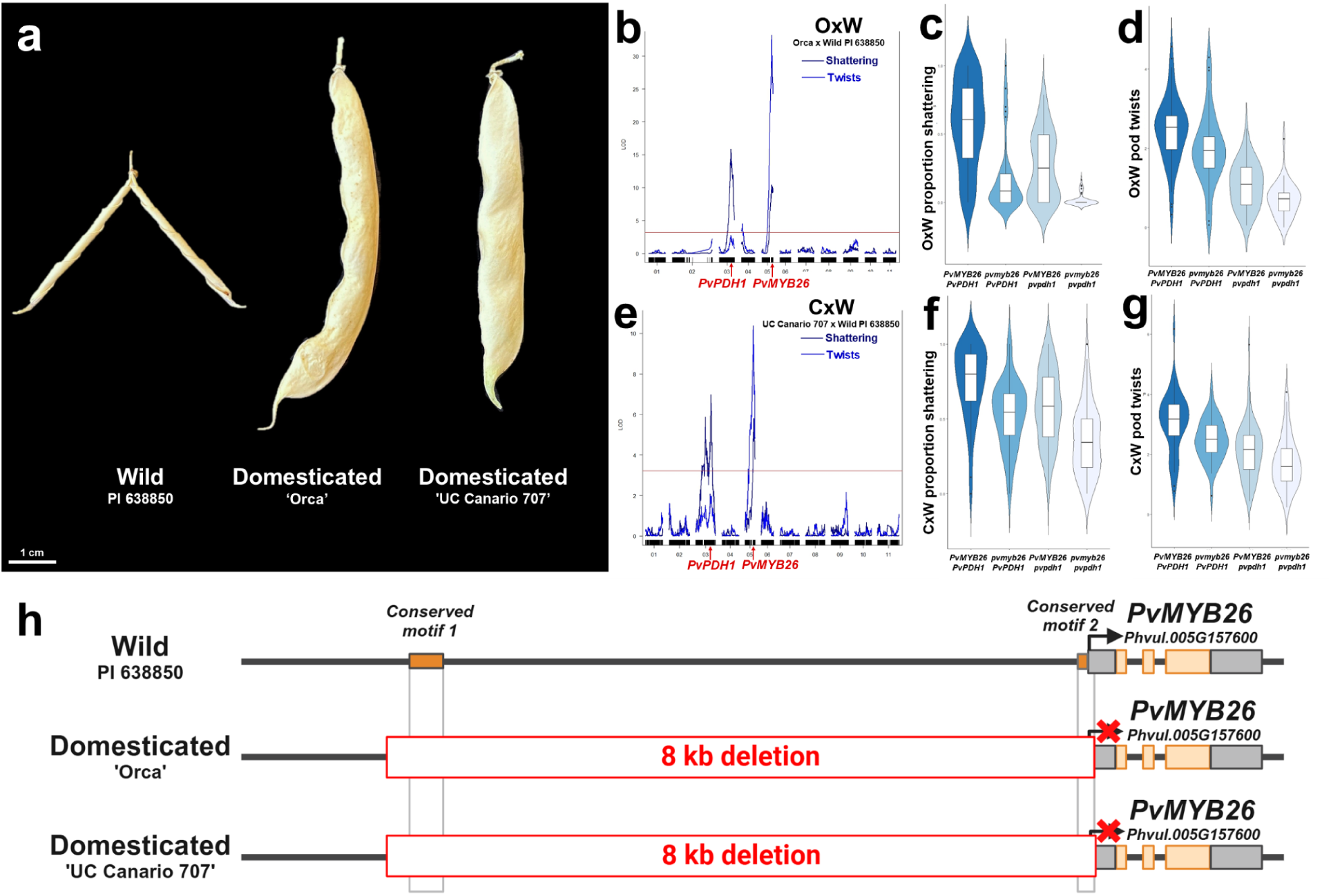
A *PvMYB26* deletion is found at a locus for reduced pod shattering and pod twists in domesticated *P. vulgaris*—a, Pod phenotypes of parental lines. The wild PI 638850 has high shattering and pod twists, while the domesticates Orca and UC Canario 707 have reduced shattering and pod twists. **b-d,** QTL mapping and allele effect plots in the Orca x PI 638850 (OxW) population identified significant QTL for pod shattering and pod twists, mapping to the candidate genes *PvPDH1* and *PvMYB26*. **e-g,** The UC Canario 707 x PI 638850 (CxW) population also had two significant QTL for pod shattering and two for pod twists, mapping to the same two candidate genes. **h,** Whole genome sequencing of the parents identified a large deletion eliminating the transcription start site, 61 bp of 5’UTR, and over 8 kb of promoter, including two motifs conserved throughout the legume family.

### A deletion eliminates 8kb of *PvMYB26*’s promoter and transcription start site in domesticated Middle American populations

Comparison of whole-genome sequencing data of the parents of the CxW and OxW populations near the pod twists and pod shattering QTL on Pv05 revealed an 8 kb deletion at *PvMYB26* in the domesticated parents Orca and UC Canario 707. The deletion, not found in the wild parent PI 638850, eliminates the transcription start site (TSS) of *PvMYB26*, including 61 bp of the 5’ UTR and more than 8 kb of promoter (Fig. 1h, Supplementary Fig. 1). *PvMYB26* was therefore considered a strong candidate to account for the reduced shattering and twisting (Fig. 2a) during the domestication of Middle American beans.

**Fig. 2.**
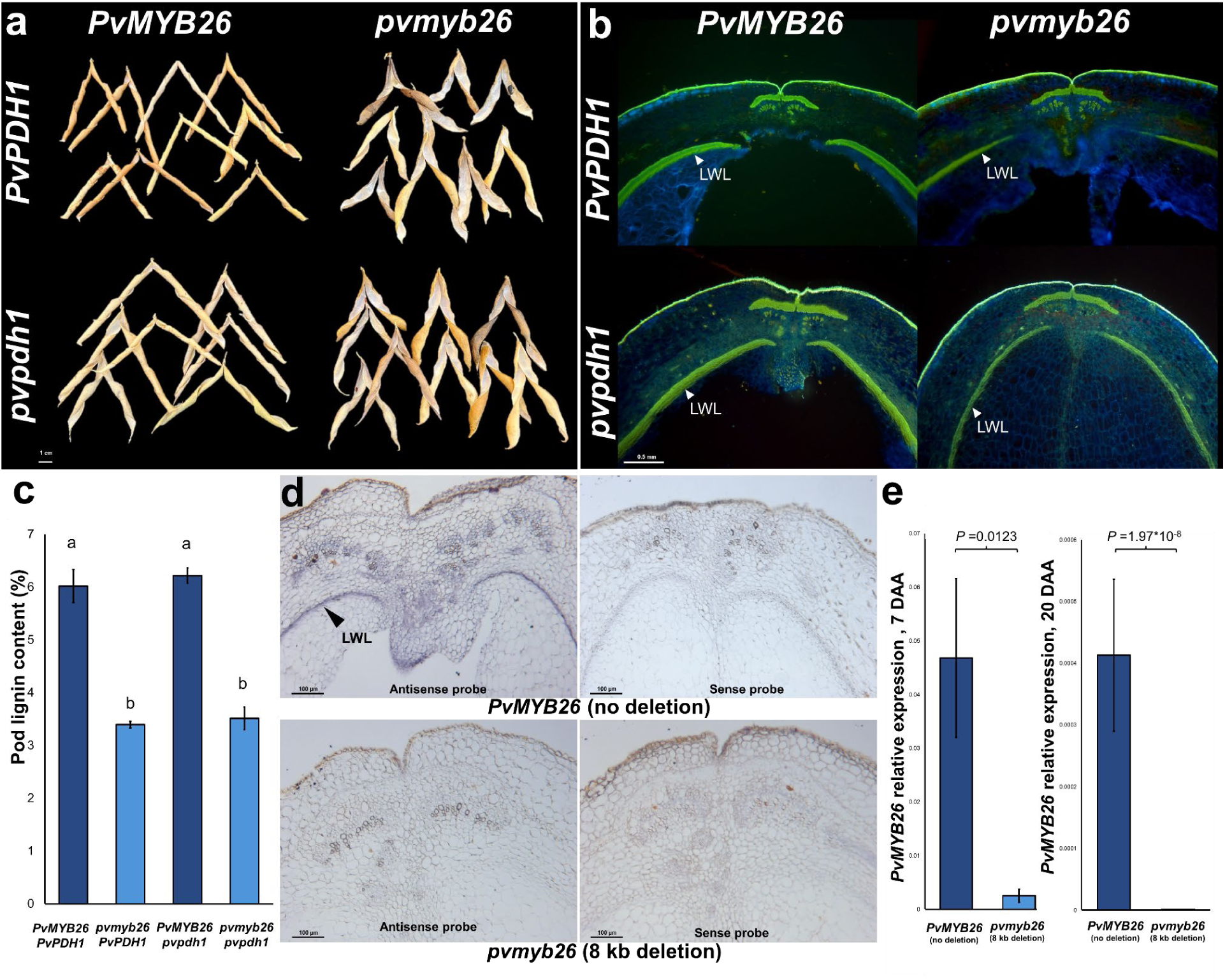
*PvMYB26* mutation effects. **a,** Visually distinct pod coiling habits in among randomly sampled pods of each allelic category, **b,** Lignified wall layer (“LWL”, hydrophobic lignin stained green) fiber deposition is visibly reduced among *pvmyb26* mutants compared to mutants, whereas *PvPDH1* has no anatomical effect, and **c,** *pvmyb26* mutants produce 44% less lignin than wild types. **d,** RNA in situ hybridization shows that *PvMYB26* expression in the pod walls is confined to the LWL, with weak additional expression in the vascular bundles. No discernible in situ hybridization signal is visible in any tissue when the 8 kb promoter deletion is present. **e,** *pvmyb26* mutants have 18.3-fold reduced *PvMYB26* expression at 7 days after anthesis (DAA) (heteroscedastic two-tailed *t*-test; *P*=0.012) and 306-fold reduced expression by 20 DAA (heteroscedastic two-tailed *t*-test; *P*=1.9*10^-8^). Data and images from OxW RILs.

To assess the functional significance of the deleted sequence through conservation analysis, we aligned *PvMYB26* and 10 kb upstream of the wild-type promoter region with 15 other legume species. Two strongly conserved noncoding sequence (CNS) motifs were identified in the *PvMYB26* promoter across all examined legume species and in both paralogous *PvMYB26* copies of each species (Supplementary Fig. 2). An 82-bp untranslated motif at the transcription start site was found in identical form in all six evaluated species of *Phaseolus* and *Vigna,* 51 bp of which was deleted in UC Canario 707 and Orca. This strong sequence conservation suggested a functional regulatory significance, warranting further analysis.

### Gene expression and lignification are dramatically reduced in types with the *PvMYB26* TSS/promoter deletion

To characterize the effects of the 8 kb *PvMYB26* TSS/promoter deletion on gene expression, qPCR was conducted on OxW RILs at two time points. At 7 days after anthesis (DAA), *PvMYB26* expression was 18.3-fold lower in *pvmyb26* mutants than in wild-type lines (Fig. 2b, *P*=0.012; heteroscedastic two-tailed *t*-test). By 20 DAA, *PvMYB26* expression was 306-fold lower in mutants (*P*=1.9*10^-8^; heteroscedastic two-tailed *t*-test), indicating that the elimination of the transcription start site and promoter had a highly significant negative impact on the gene’s transcription level. Additionally, analysis of published pod RNA-seq data^32^ of four accessions across three pod sampling time points showed that *PvMYB26* expression was significantly decreased in mutant compared to wild-type individuals (*P* = 5.904*10^-5^, linear mixed model, Supplementary Fig. 3).

Microscopy and pod biochemical analyses were then conducted to determine the anatomical and biochemical effects of the *PvMYB26* deletion. *PvMYB26* mutants showed greatly reduced lignin deposition in the lignified wall layer (LWL) of pods, without observable differences in the bundle sheath, dehiscence zone, or other structures (Fig. 2c). Wild-type pods were composed of 6.0% lignin by mass. In contrast, mutant pods contained only 3.4% lignin, representing a statistically significant decrease of 43.5% in mutants (protected Fisher’s LSD, Fig. 2d). In contrast, *PvPDH1* allele had no significant effect on pod lignin content, confirming that its effects on pod shattering are independent of lignin quantity^33^.

To further investigate the spatial expression pattern of *PvMYB26*, we performed mRNA in situ hybridization on pod sections sampled seven days after anthesis (DAA). Strong hybridization signals were specifically detected in the LWL cells of the pod walls (Fig. 2d), with minor expression observed in adjacent bundle sheath tissues. No visualizable hybridization signal was identified when the 8 kb deletion was present (Fig. 2d). These results indicate that *PvMYB26* expression occurs in the LWL, which is known to influence pod torsion and is differentially thickened between mutants and non-mutants^5^.

### A 125 kb domestication-related selective sweep occurred at *PvMYB26*

An initial assessment of the *PvMYB26* TSS/promoter deletion frequency in domesticated and wild materials was conducted by designing a high-throughput co-dominant PCR assay, which was screened on a collection of 300 *P. vulgaris* accessions. Strikingly, the deletion was found in 99.1% of Middle American domesticates, but just 2.2% of wild beans (*F_ST_* = 0.94). The deletion, which is Middle American in origin, had also been introduced into 14% of accessions of the independently domesticated Andean Diversity Panel (Supplementary Table 2, Supplementary Fig. 4).

To better understand the haplotype diversity of PvMYB26 and the geographic origin of the eight-kb mutation, whole-genome sequencing and analysis were then conducted on 326 lines. Only one among 55 Middle American wild lines (1.8%; PI 201016) showed the deletion. Of 138 domesticates with a Middle American *PvMYB26* haplotype, the deletion was found in 97.1%, only absent in four Mexican landraces (Fig. 3b). *F_ST_* for the deletion was 0.89 in this panel.

**Fig. 3.**
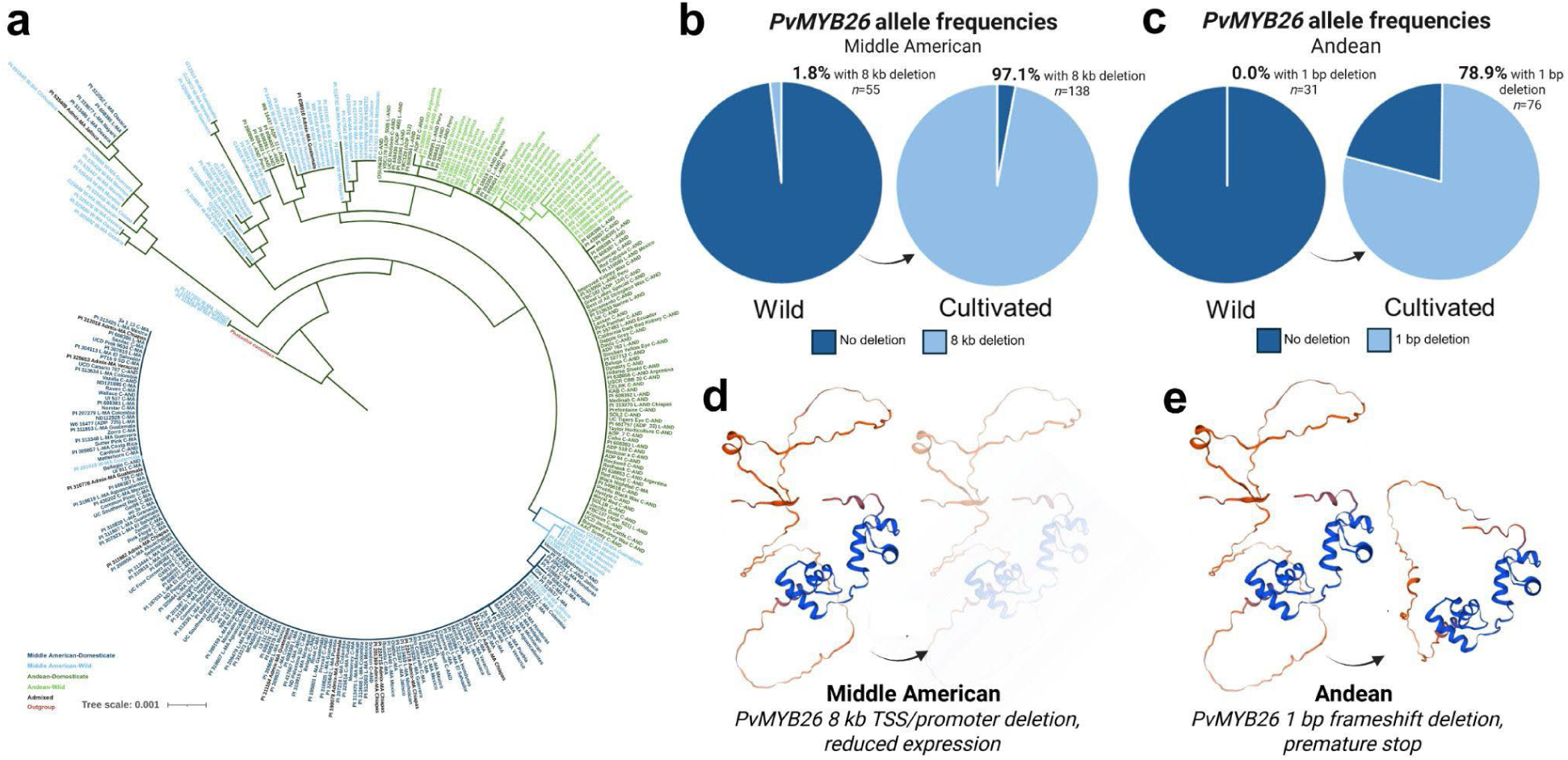
Sequence diversity at *PvMYB26* among domesticated and wild populations. **a,** A dendrogram of sequence diversity based on 4 kb of sequence at *PvMYB26*, colored by accession gene pool and domestication status. **b,** The 8 kb deletion is found in only 1.8% of sampled Middle American wild types (PI 201016, with admixture and large seeds). In contrast, the mutation is found in 97.1% of surveyed cultivated lines with a Middle American haplotype. **c,** A distinct 1 bp mutation arose among Andean domesticates, which was not found in any wild type but was present in 78.9% of domesticated lines with an Andean haplotype. **d,** Whereas the Middle American 8 kb TSS/promoter deletion drastically reduces gene expression, **e,** the Andean one-bp mutation disrupts the final 79 amino acid chain and truncates its length by 45 residues. The extreme delineation between wild and cultivated types at *PvMYB26* indicates that mutations at the gene likely have differential fitness for plants in wild vs. cultivated ecosystems, and are not readily exchanged between wild and domesticated populations.

To investigate the possibility of a broader selective sweep, we calculated Tajima’s D, nucleotide diversity, and *F_ST_* over a 600 kb interval centered on *PvMYB26*. Using all three metrics, a hard selective sweep was identified in the Middle American gene pool over 125 kb (chromosome Pv05 43.977 Mb to 44.102 Mb; 5_593 genome v1.1). The 125 kb region exhibited a mean Tajima’s D of -2.05 ± 0.03 in domesticates, whereas the average value outside this sweep interval was 1.16 ± 0.10 (Fig. 4a, Supplementary Table 3). Tajima’s D showed no significant reduction in the region among wild lines (Fig. 4d, Supplementary Table 3). In the sweep interval, domesticates had just 12% ± 0.7% of the nucleotide diversity found in wild types, whereas in the surrounding region they had 53.4% ± 1.7% of the diversity found in wild types (Fig. 4b,c). An increased mean *F_ST_* of 0.49 ± 0.01 was also observed between wild and domesticated lines, compared to a mean *F_ST_* of 0.25 ± 0.01 in the surrounding region (Fig. 4d). All of these statistics indicate that a hard sweep occurred across the region among domesticated Middle American populations, differentiating it from wild types.

**Fig. 4.**
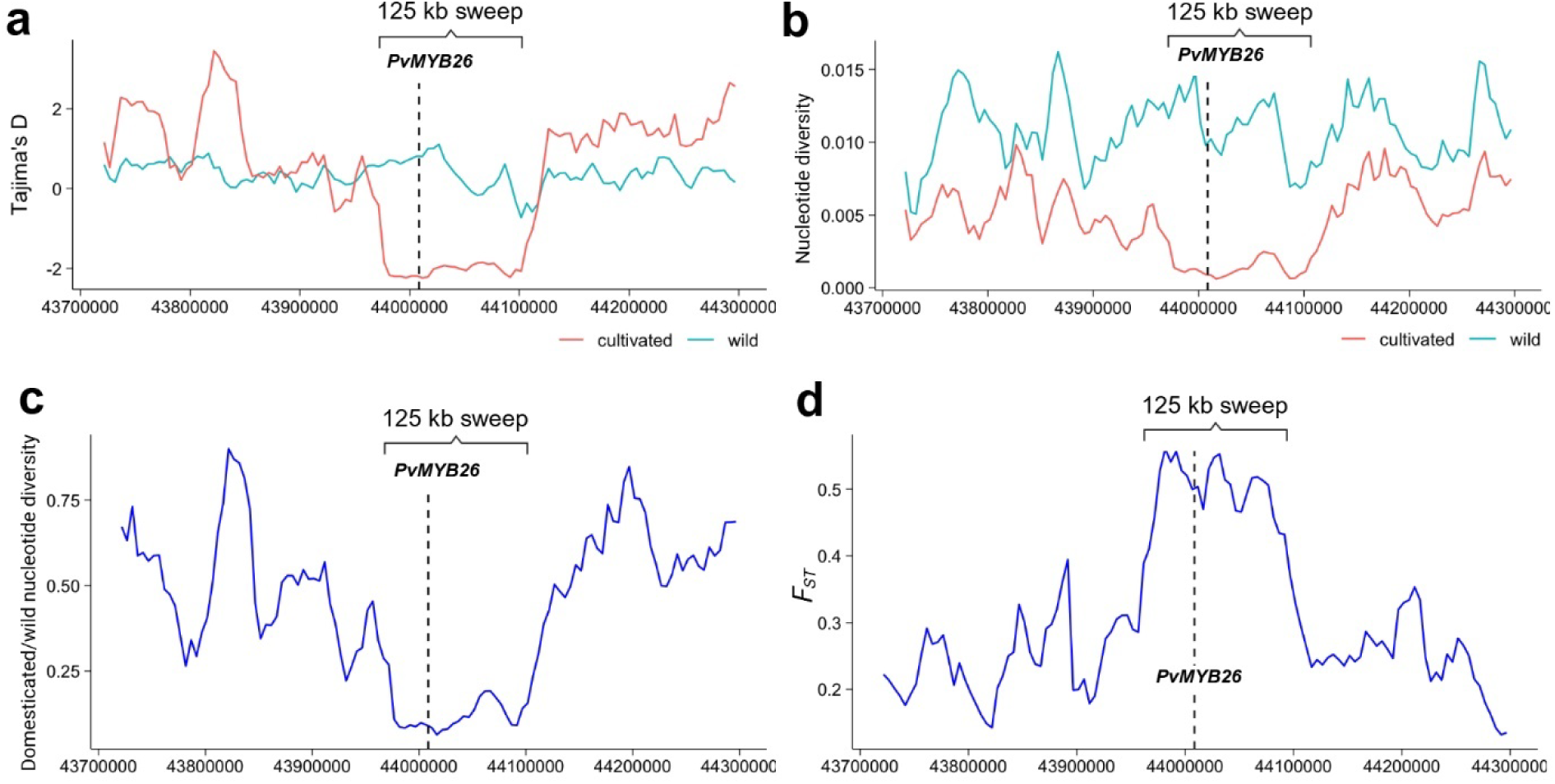
A 125 kb selection sweep occurred at *PvMYB26* during the domestication of Middle American common beans. **a,** Tajima’s D in wild and domesticated populations shows that values drop around *PvMYB26* among domesticates, while wild type values are stable, **b,** and **c,** Wild lines have stable nucleotide diversity in the region, while nucleotide diversity drops around *PvMYB26* among domesticates, **d,** *F_ST_* between wild and domesticated populations is highest in the selective sweep region, showing high distinctiveness between populations. Coordinates relative to genome 5-593 v1.1.

### A 1 bp *PvMYB26* frameshift deletion exists solely in Andean domesticates

The whole-genome sequencing data were also analyzed for variation that might be responsible for reduced shattering in the independently domesticated Andean gene pool of common bean. We identified a one-bp frameshift deletion in exon 3 of *PvMYB26* (38,639,348 bp), leading to a complete disruption of the remaining 79 amino acid sequence and introducing a premature stop codon, which truncates its length by 45 amino acids. The deletion is found in 78.9% of Andean domesticates (*n*=76) but is absent among wild beans (*F_ST_* = 0.66).

Seven wild lines (PI 638850, PI 390770, PI 638864, PI 638872, W6 18809, W6 18810, W6 18811) lacking the 1 bp deletion had otherwise identical haplotypes to this mutant sequence. These wild types spanned three countries (southern Peru, Bolivia, and Argentina), and no selection sweep was identified in the genome area around *PvMYB26* (Supplementary Table 4, Supplementary Fig. 5). The 1 bp deletion was therefore not considered informative in identifying the specific geography of domestication in Andean beans.

### West-Central Mexico as a center of origin for Middle American agriculture

The extreme *F_ST_* and near total lack of gene flow between wilds and domesticates at *PvMYB26* offered a unique opportunity to identify the geographic origins of Middle American beans without the effects of historical admixture. To this end, consensus FASTA sequences within 2 kb of the *PvMYB26* promoter deletion were extracted and aligned with the predominant domesticated haplotype. The wild haplotypes most similar to the domesticated haplotypes were found in eastern Jalisco (PI 417707, PI 417708). The entire coding DNA sequence of *PvMYB26* is identical to that of domesticates in these types, with only a one-bp promoter indel distinguishing them from the predominant domesticated haplotype. Accessions from northern Michoacan (G12894, PI 417621) also had *PvMYB26* transcribed sequences identical to domesticates. Three wild accessions from Morelos (PI 325679, PI 535430, PI 535444) and one from southern Guanajuato (G23433) had promoter sequences identical to domesticates but differences in the transcribed region. The identification of wild haplotypes most similar to domesticates in this region suggests that this area was the site of origin for the *pvmyb26* mutation that strongly distinguishes wild and domesticated Middle American populations, and indicates the region was likely the Middle American center of domestication of common bean (Fig. 5).

**Fig. 5.**
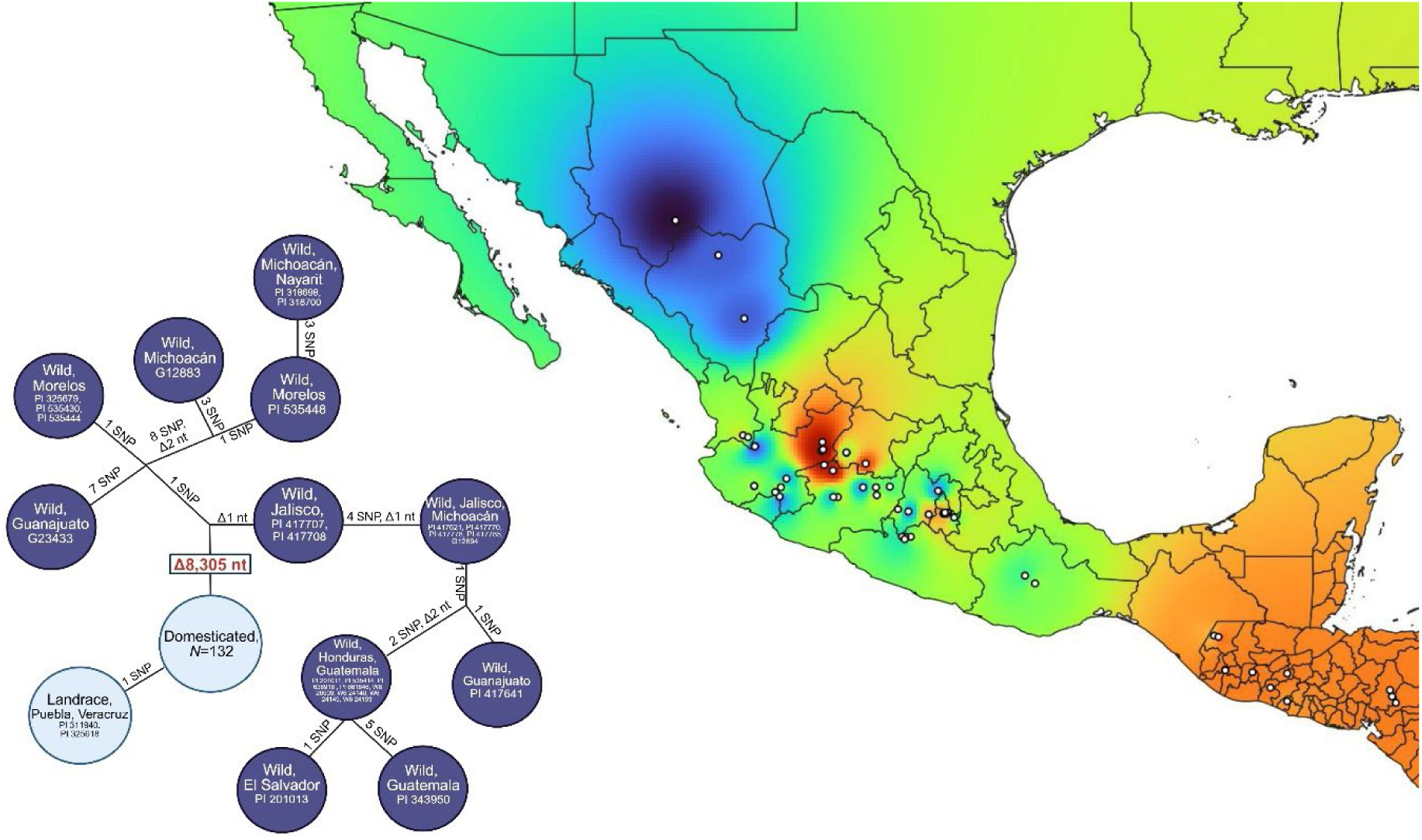
The wild form of *PvMYB26* most like the domesticated haplotype is found in eastern Jalisco and northern Michoacán, in west-central Mexico, indicating a likely domestication event in the region. As the common bean is a central part of Mesoamerica’s *milpa* or “three sisters” agriculture (along with maize and squash), these results shed light on the geographic origins of this important form of New World agriculture and food production. Warmer map colors indicate greater similarity to domesticates. Haplotype network colors indicate deletion status.

## Discussion

In grain-propagated crops, including cereals, pulses, and *Brassica* oil crops, the absence of grain release at maturity is of utmost importance to maximize yields under cultivation. Preventing grain shedding has taken multiple anatomical appearances, including acquiring a stiff rachis through the disruption of abscission layer formation in Fertile Crescent cereals^4,34^ and losing explosive pod shattering in grain legumes, due to reduced pod torsion force or changes in dehiscence zones^5,9^. Identifying the molecular basis of crucial domestication traits sheds light on the entire domestication process and more broadly on the origins of agriculture^2,35^. Here, we document research results leading us to the identification of a single major gene – *PvMYB26* – which likely reduced pod shattering in the initial stages of the two domestications of common bean, in Middle America and the southern Andes^17,36^. Furthermore, a comparison of haplotypes in and around the locus suggests a domestication area for the Middle American common bean and, more broadly, an initiation of Mesoamerican agriculture in west-central Mexico.

The evidence supporting a key role of the *PvMYB26* gene on chromosome Pv03 in pod shattering includes genetic linkage mapping in two recombinant inbred populations between a domesticated and a wild bean parent and the identification of an eight-kb deletion affecting the upstream promoter-containing sequence and the transcription start site in Middle American domesticates. This deletion mutation strongly reduces the gene’s expression, is correlated with reduced lignin deposition in the pod walls and is accompanied by a substantial reduction in genetic diversity in nearly every Middle American domesticated type tested (*n* = 134) compared to the corresponding wild progenitor, characterizing a marked selective sweep of some 125 kb in the genome region in and around the gene. These sequence features indicate a Middle American origin in west-central Mexico for the mutation, with subsequent introgression in a minority of the Andean domesticated gene pool. In contrast, most of the Andean domesticated gene pool shows a one-bp frameshift deletion in *PvMYB26*, causing a truncation and loss-of-function of the gene product. All these results strongly suggest a primordial role for *PvMYB26* in pod dehiscence and bean domestication.

The *PvMYB26* hard sweep, extending over 125 kb, is significantly larger than the 66 kb selective sweep found at the *sh4* shattering locus in rice^37^, or the 65.6 kb sweep at the *tb1*^38^ in maize, both of which are key domestication genes in their respective species. The size difference may be attributed to stronger linkage disequilibrium in common bean compared to the two cereal species and the limited gene exchange between wild and domesticated beans, especially around domestication genes^26-28^. The hard sweep around *PvMYB26* contrasts with the multiple mutations and soft sweeps that have been commonly observed among domestication syndrome traits across species^39,40^, including common bean^41-45^. In the Andean gene pool, where the selective sweep at *PvMYB26* was weaker (Fig. 4C, Supplementary Table S4, Supplementary Fig. S5), a greater number of secondary loci have been described^33^. There has also been a significant introgression of the Middle American mutant haplotype into the Andean gene pool (Supplementary Table S2, Supplementary Fig. S4).

The expression pattern of *PvMYB26*, which was limited to the LWL fibers, correlates with the observed lignin deficiency phenotype in *pvmyb26* mutants and supports *PvMYB26’s* direct role in regulating lignification within this specialized cell layer. The connection between MYB transcription factors and lignin biosynthesis is well-established across plant species^46-52^. In Arabidopsis, *MYB58* and *MYB63* directly activate lignin biosynthetic genes like *PHENYLALANINE AMMONIA LYASE 1* (*PAL1*), *CINNAMATE 4-HYDROXYLASE* (*C4H*), and *4-COUMARATE COA LIGASE 1* (*4CL1*), by binding AC-rich promoter elements^47,53^. Similarly, *MYB26* regulates secondary cell wall thickening in anthers by controlling *NAC SECONDARY WALL THICKENING PROMOTING FACTOR1* (*NST1*) and *NST2* expression, while *MYB46*/*MYB83* orchestrate a hierarchical network activating both lignin biosynthesis and secondary wall formation^46,47^. These findings align with our observations that *PvMYB26* disruption specifically impacts LWL lignification, highlighting the precision of MYB-mediated transcriptional control in spatial regulation of lignification. The conserved role of MYB transcription factors in activating lignin pathway genes provides a mechanistic context for the severe lignin reduction caused by the *PvMYB26* promoter deletion.

While *PvMYB26* may have been sufficient for reduced pod shattering during the initial domestication phase of common bean, it was not sufficient for control of pod shattering in arid environments, which exacerbate pod shattering. In these environments, additional mutations in the *PvPDH1* dirigent gene on chromosome Pv03^5,33,54^ and other genes, selected after domestication, provided additional resistance to pod shattering and local ecogeographic adaptation (Fig. 6). *PvMYB26* had previously been described as an improvement gene selected subsequently in the development of modern snap beans^7,21,22^. However, the near-uniform presence of the 8-kb mutation in the domesticated Middle American gene pool strongly suggests that PvMYB26 is a foundational component of the initial domestication process of Middle American common beans. Identifying the one-bp frameshift mutation in 79% of dry beans with Andean haplotypes also indicates its importance in the initial domestication of that gene pool.

**Fig. 6.**
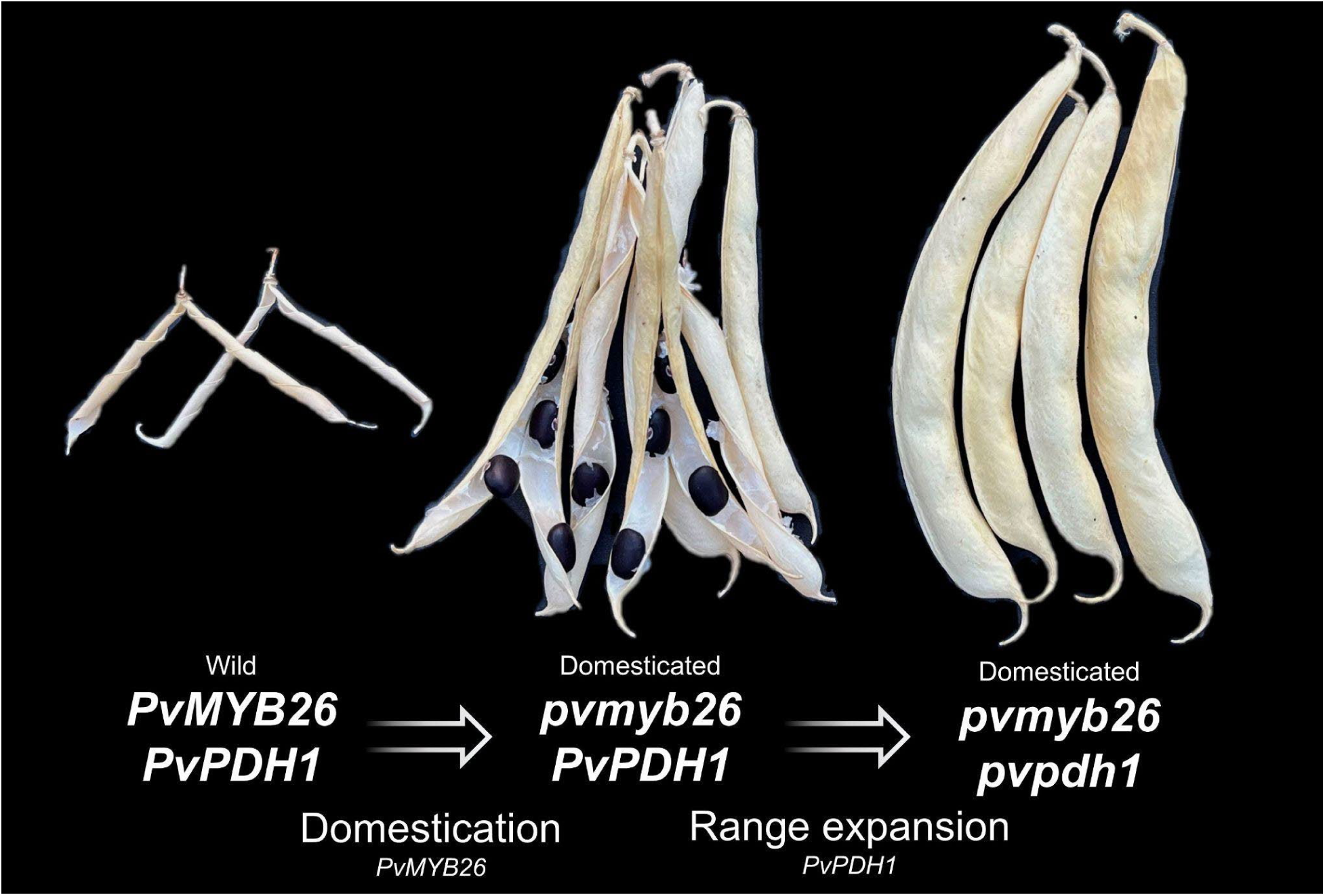
Overview of pod changes during the domestication and range expansion of the Middle American common bean. During domestication, a mutation in *PvMYB26* on Pv05 led to a reduction in the quantity of wall fiber between wild and domesticated Middle American beans with intermediate shattering. Varieties of the ecogeographic race Durango are adapted to arid conditions where pod shattering is most strongly expressed and have accumulated a mutation in *PvPDH1* on chromosome Pv03^33^.

Our results further suggest that, at least in the case of pod shattering, orthologous genes may have been involved across grain legume domestications. In both common bean domestications – Middle American and southern Andean – mutations in the same gene that reduced pod shattering were selected. This observation raises the question of whether other, more distantly related grain legumes show a similar convergence. The legume family was domesticated approximately 40 times independently, more than any other plant family^5^, creating an unequaled natural experiment in parallel evolution. Growing evidence indicates that *MYB26* homologs are critical regulators of seed dispersal and carpel development across legumes, contrasting their anther-specific role in Arabidopsis^46,55^. In *Vigna* sp., loss of pod shattering has been linked to loss-of-function mutations in *MYB26* orthologs in five independently domesticated lineages^56,57^. These *Vigna MYB26* mutants also show reduced fiber deposition in the pod LWL, consistent with our anatomical and biochemical results and other studies in common bean^21,58^. Unlike some *Vigna* mutants, however, the TSS/promoter deletion of *PvMYB26* does not lead to a complete elimination of pod lignin in the LWL, which may be advantageous for threshing during harvest as it facilitates the recovery of grains with limited damage.

Similarities in protein or nucleic acid sequences^2,35,59-61^ between wild and domesticated types of crop species can provide information about the geography of domestication, including the number and location of putative domestication regions, with the caveat that introgression can have a confounding effect in this geographic assessment. In this case, the almost exclusive presence of the mutant *PvMYB26* haplotype in the domesticated Middle American gene pool and its near absence in the sympatric wild gene pool confirmed the restricted gene flow around domestication genes observed by Papa et al.^27,28^ and limited any confounding effects due to admixture. The comparison of the two-kb haplotype surrounding the *PvMYB26* deletion showed that wild haplotypes most like the domesticated ones were situated in a region encompassing eastern Jalisco and northern Michoacán (Fig. 5). This region is largely congruent with a region identified by Gepts^23^ and Kwak et al.^24^, using different genome-wide molecular markers. This region overlaps the current area and includes areas to the east in Guanajuato and western Jalisco in the Rio Lerma-Rio Santiago basin.

It is striking that this area is overlaps with a region in which the presumed ancestors of the three major crops of the principal cropping system (milpa system) are distributed: beans (*Phaseolus* spp.), maize (*Zea mays* L.), and squash (*Cucurbita* spp., and more specifically *C. argyrosperma*) (Zizumbo-Villarreal and Colunga García-Marín 2010). A putative domestication region for maize and *C. argyrosperma* has been widely hypothesized to be in the lower Balsas River basin^62-66^, leading Zizumbo-Villarreal and Colunga García-Marín^67^ to suggest that the northwestern part of the Balsas-Jalisco biogeographic region, which encompasses the two largest river basins of Mexico, could have witnessed the initiation of the cultivation and domestication of the three major component crops of the milpa agroecological food system and, hence, agriculture in Mesoamerica. Further cultural developments led to the expansion of the basic milpa cropping system by integrating additional crops and animals and developing more complex food systems supporting the diverse pre-Hispanic civilizations^68,69^.

Finally, the following evolutionary pod domestication scenarios can be proposed. In the Middle American gene pool (Fig. 6), more specifically in what is now west-central Mexico (Jalisco, northern Michoacan, and western Guanajuato), selection for reduced pod shattering during the initial phase of domestication may have led to the *de novo* selection and near-fixation of an 8-kb deletion affecting the *PvMYB26* locus on chromosome Pv05 (current results). In a subsequent phase, differentiation of the incipient MA gene pool into ecogeographic races and dissemination across the diverse ecosystems of the Middle American area^36,70^, selection for nearly complete pod indehiscence in the arid northern Mexican highlands led to the preponderance of a *PvPDH1* (Pv03) loss-of-function, non-synonymous substitution in race Durango, which includes bean types such as pintos, pinks, and great northerns^54^. In contrast, a separate migration of Middle American domesticates towards the Middle American lowlands (race Mesoamerica, represented by smaller-seeded navy and black beans^70^) did not result in selection for additional pod shattering tolerance^54^, given the more humid nature of the habitat. A similar scenario, involving the same genes, can be developed for the Andean domestication. Reductions in pod shattering can be traced to either the one-bp mutation in *PvMYB26*, to the introgression of the 8-kb mutation from Middle American domesticates, or selection of hitherto undetermined pod shattering genes in the Andean gene pool. As observed earlier, however, the reduced diversity of the latter gene pool prevents us from identifying a more specific location of the Andean domestication anywhere between southern Peru and northwestern Argentina.

Understanding the genetic basis of domestication enhances the ability to identify critical genetic diversity in landraces and crop wild relatives maintained in gene banks to develop more resilient crops and improve global food security.

## Online methods

### Plant Materials and Phenotyping

‘Orca’ (domesticated Middle American common beans^71^) and ‘PI 638850’ (wild Andean common bean) were used as parents to develop a 292-member “OxW” RIL population segregating for domestication-related traits, which were phenotyped at the F_4_ and F_5_ generations. The domesticated Andean bean ‘UC Canario 707’^72^ and wild PI 638850 were used as parents for a second RIL population, (“CxW”), consisting of 269 members phenotyped at the F_5_ and F_6_ generations. While UC Canario 707 is primarily Andean in ancestry, it contains a Middle American haplotype at *PvMYB26*. Orca and UC Canario 707 are highly resistant to shattering, while PI638850 is highly prone to shattering, typical of wild beans. RIL populations were planted in the University of California, Davis greenhouse during the fall and winter of 2022-2023 and 2023-2024, to allow for the long nights required for flowering in photoperiod-sensitive wild materials. Temperatures were maintained between 15°C at night and 26°C during the day. Dry pods were harvested before shattering and allowed to dry for at least 30 days at room temperature in laboratory conditions to express pod shattering. The proportion of pods shattering was then quantified. To test pod twists, non-shattered pods were then artificially split and then desiccated through incubation at 40°C for two weeks, with an additional two weeks of re-equilibration at room temperature. The number of pod twists (rotations) was then taken from three representative pods of each RIL.

Initial marker screens were run on a panel of 300 lines: 90 wild types, 110 Middle American Diversity Panel (MDP)^73^ domesticates, and 100 Andean Diversity Panel (ADP)^74^ domesticates. This panel of 300 lines included all wild accessions in the NPGS gene bank with seed sizes smaller than 15 g/100. Domesticates were chosen randomly among the diversity panels.

Whole-genome shotgun sequencing was conducted on a panel of 289 novel lines, analyzed along with 37 recently published genomes^45^. In total, the collection included 326 lines, of which 225 were domesticated, 89 were wild, and 12 were admixed. According to genome-wide sequencing data, accessions were classified as domesticated, wild, or admixed.

### Genetic mapping

Genotyping of the CxW and OxW populations was conducted according to the Khufu method^75^ by the HudsonAlpha Institute (Huntsville, AL, USA), yielding 48,197 initial SNPs in the CxW population and 189,595 initial SNPs in the OxW population. SNP data were cleaned using the methods of Xie et al.^76^. After filtering co-segregating markers, 2,113 SNPs were retained in the CxW population, and 2,156 were retained in the OxW population. The ASMAP and R/QTL packages were used for linkage and QTL mapping^77-79^ using the Haley-Knott algorithm^80^. The fitqtl() function was used to assess allelic interactions and effects. A LOD significance threshold of 3.21 was used based on the 95% threshold of 1000 randomized permutations of the data. Allele effect plots were visualized in ggplot2^81^.

### Sequencing and analysis

Illumina 150 bp paired-end sequencing was conducted to a depth of 12-16x was conducted on the 289 novel accessions at the UC Davis Genome Center’s DNA Technologies Core and the HudsonAlpha Institute for Biotechnology (Huntsville, AL, USA). DNA for sequencing was isolated using the Qiagen DNeasy plant mini kit, verified for quality by Nanodrop and gel. Raw sequences were quality-checked with FASTQC^82^ and trimmed using Trimmomatic v0.39’s ILLUMINACLIP function^83^. The Burrow-Wheeler Aligner (BWA-MEM^84^) was used with default settings to align trimmed reads to the reference genomes G19833 v2.1^19^ (https://phytozome-next.jgi.doe.gov/info/Pvulgaris_v2_1) and 5-593 (https://phytozome-next.jgi.doe.gov/info/Pvulgaris5_593_v1_1). Consensus sequence was extracted using samtools consensus^85^. Alignments were visualized in the Integrative Genomics Viewer (IGV)^86^. Protein models were generated using the SWISS-MODEL Expasy web server [SWISS-MODEL (expasy.org)].

To identify conserved non-coding sequence, *PvMYB26* and 10 kb of its promoter sequence were aligned with 15 other legume species extracted from Phytozome 13^87^ (accessed 14 September 2024), as well as the *Pisum sativum* assembly cultivar ‘Zhewan’ from NCBI (https://www.ncbi.nlm.nih.gov/datasets/taxonomy/3888/, accessed 24 January, 2025), with sequence alignment conducted using NCBI BLASTn.

### Microscopy

Pods from the OxW population were collected at stage R8 (full-seeded green pod) from lines characterized at allele for both *PvPdh1* and *PvMYB26* genes based on selectable markers for each gene (Supplementary Table 5). Sections 100 μm thick were made using a vibratome and treated 0.01% Auramine O and 0.007% Calcofluor solution for 20 minutes^88^. These two stains enable the secondary cell wall components in plant tissues to be distinguished from each other under the microscope, with Calcofluor dying cellulose blue and Auramine O staining lignin and other hydrophobic tissues (e.g., cuticle) green. Images were visualized using a fluorescence microscope (Olympus BH2-RFL, Waltham, MA, USA) with ultraviolet filter set (UG-1 and DM-400 + L-420).

### mRNA *in situ* hybridization

mRNA *in situ* hybridizations and probe synthesis were performed as previously described^89,90^. Briefly, bean pods of six OxW RILs were collected at seven days after anthesis (DAA). Pods were hand-sectioned transversely and fixed in 4% paraformaldehyde (Electron Microscopy Sciences). Fixed tissue was dehydrated through a graded ethanol series (50%, 70%, 85%, 95%, and 100%), followed by a Histoclear series, and embedded in paraffin (epredia Precision Cut, B1002490). Sections of 10 μm thickness were cut using a Leica microtome and mounted on ProbeOn Plus Slides (Fisher Scientific, 22-230-900). Primer sequences used to amplify *PvMYB26* for probe synthesis were TACACACCAGGCATTCGTGC and GAACTAGCATGGCTCTGAGGT. Images were acquired using an OLYMPUS BX50 DIC microscope.

### Expression analyses

Pod samples were harvested at two growth stages, 7 and 20 days after anthesis (DAA). Pod sections 1 cm thick were taken with a razor blade, and seeds were removed from the 20 DAA samples sections. Samples were flash frozen in liquid N_2_ and stored frozen at -80°C. Pod sections were ground in liquid N_2_ using a mortar and pestle, and total RNA was extracted using the RNeasy Plant Mini kit (Qiagen, Hilden, Germany). Concentration and quality were subsequently confirmed by NanoDrop and 1% bleach gel. Then 3 µl RNA per sample was converted to cDNA using SuperScript IV VILO Master Mix with ezDNase. RT-qPCR primers were designed using NCBI primer blast. *Act11* (*Phvul.002G269900*) was used as a control gene. Primers spanning intron-exon junctions were used for both genes (Supplementary Table 1). qPCR primer efficiency was checked on pooled cDNA from each sample. For 7 DAA, *n*=11 for *PvMYB26/PvMYB26*, *n*=7 for *pvmyb26/pvmyb26*; for 20 DAA *n*=11 *PvMYB26/PvMYB26*, *n*=5 for *pvmyb26/pvmyb26*. For ΔC_T_ calculation, the C_T_ values of the reference gene were subtracted from the C_T_ values of the candidate gene. These values were then converted to 2^-ΔCT^ values. The mean, standard deviation, and standard errors of 2^-ΔCT^ data for both *PvMYB26* wild and domesticated allele lines were calculated. Expression differences between these two alleles were compared using heteroscedastic two-tailed *t*-test.

Pod RNA-seq data from Parker et al.^32^ were analyzed for four genotypes: Midas, G12873, ICA Bunsi, and SXB 405; at three different stages (R7, pod formation; R8, pod fill; and R9, pod maturation^91^). Three replicates were conducted for each stage and genotype. The expression patterns of ICA Bunsi and SXB 405 were compared with the expression patterns of Midas and G12873 to identify transcripts differentially expressed between *PvMYB26* mutants and wild types. The expression of candidate loci was compared by mixed linear model with mutation status as a fixed effect and maturity stage as a random effect.

### Lignin content

OxW RILs were classified according to genotype at both *PvMYB26* and *PvPDH1* using codominant PCR markers, with four total allelic categories. For each RIL and year, three pods were assigned to one of four technical replicate bins for each of the four corresponding allelic categories. These dry pods were then submitted to Cumberland Valley Analytical Services for lignin quantification. Lignin mass was determined using the Goering and Van Soest^92^ method with modifications. The fiber residue from the acid-detergent fiber (ADF) extraction was recovered on a Whatman Glass Fiber Filter in a California Buchner Funnel. Approximately 45 ml of 72% Sulfuric Acid was added to the fiber residue, followed by shaking for two hours to ensure that fibers were washed with acid. After the acid-washed fiber residue was filtered, rinsed, dried and weighed, lignin residue was washed for two hours to remove lignin organic matter. The ash residue was weighed again, and the amount of lignin was determined by subtracting it from the original weight. Results were analyzed using an ANOVA-protected Fisher’s LSD test based on mutation status at *PvMYB26* and *PvPDH1*.

### Haplotype diversity

To generate haplotype networks and gene trees, consensus FASTA within 2 kb of the *PvMYB26* TSS/promoter deletion was extracted from the reference genome aligned whole genome sequencing lines using samtools consensus function^85^. This 4 kb region included the full transcribed region of the gene and approximately 2 kb of promoter. Alignments were built with MAFFT (version 7)^93^. A single gene tree was estimated with IQ-TREE (v.1.6.12)^94^ under the GTR+G model^95^ with 1,000 bootstrap replicates. The phylogenetic tree was visualized by the iTOL online server^96^. To find out the haplotype network, the NeighborNet algorithm was implemented in the SplitsTree program (version 6.4.11)^97^.

Population genetics statistics were calculated on biallelic variants identified within a 600 kb region centered on *PvMYB26*. A vcf was generated from .bam files using bcftools mpileup and bcftools call, based on the 5_593 v1.1 reference genome. Principal component analysis was performed using PLINK 1.9^98^ and used to classify materials as having Andean, Middle American, or admixed haplotypes at the region. Wild, domesticated, and admixed status were classified based on genome-wide SNP data, identifying several misclassified lines in the NPGS-GRIN database. The biologically distinct and independently domesticated Middle American (wild *n*=55, domesticated *n*=139) and Andean (wild *n*=31, domesticated *n*=76) gene pool materials were then processed and handled separately for subsequent steps. The R package PopGenome^99^ was used to calculate nucleotide diversity, *F_ST_*, and Tajima’s D. Variants were loaded using the option “include.unknown=TRUE” after determining that the level of missing data was not substantial enough to bias the analyses when run in nucleotide mode. A sliding window transform was used to split the variants into 25 kb windows with a 5 kb step. Statistics were plotted using the R package ggplot2^81^ with the position on the x-axis representing the position at the center of each window.

### Geographic Analysis

Heatmaps of similarity between wild lines and the core domesticated haplotype were generated using the consensus sequences within 2 kb of the TSS/promoter deletion used for haplotype network analysis. Consensus sequences were aligned to the predominant Middle American domesticated haplotype of accession 5-593 using NCBI BLASTn. E scores and geographic origins of wild Middle American accessions were then loaded into QGIS v3.22.6. Heatmaps were generated using the IDW interpolation function with a distance coefficient set to *P*=3. National, state, and oceanic boundaries were downloaded from Natural Earth data (Kelso and Patterson 2010).

## Acknowledgements

This project was made possible by germplasm from the United States Department of Agriculture National Plant Germplasm System (Pullman, Washington, U.S.A.) and the gene bank of the International Center for Tropical Agriculture (CIAT, Cali, Colombia). We thank Daniel Debouck for important germplasm advice. Judy Jernstedt provided microscopy equipment. Maizie Flores, Anham Rafique, and Troy Williams assisted in population development and phenotyping. This work was funded by Kirkhouse Trust A23-1518-001 to T.P. and USDA-AFRI award 2023-67013-40001 to T.P. X.X. acknowledges support from the University of California, Davis (UC Davis) new faculty start-up funds and California Agricultural Experiment Station/National Institute of Food and Agriculture (CA-D-PLB-2850-H).

## Author contributions

T.P., P.M., and P.G. conceived the research; T.P., B.C., and P.M. designed the experiments; T.P., P.M., and A.F. acquired funding; B.C., T.P., J.R., A.F., S.E., X.Y., X.X., and P.M. performed the analyses. B.C. and T.P. drafted the first version of the paper. All authors contributed to discussions and revisions of the paper.

## Data deposition

Sequences aligning within 300 kb of *PvMYB26* and used for population genetics screens will be submitted to the NCBI SRA. Mapping data will be submitted to Data Dryad. Other published datasets used in this study are listed in the related legends, Online Methods, and/or supplement.

## Supplementary Tables

**Supplementary Table 1.**
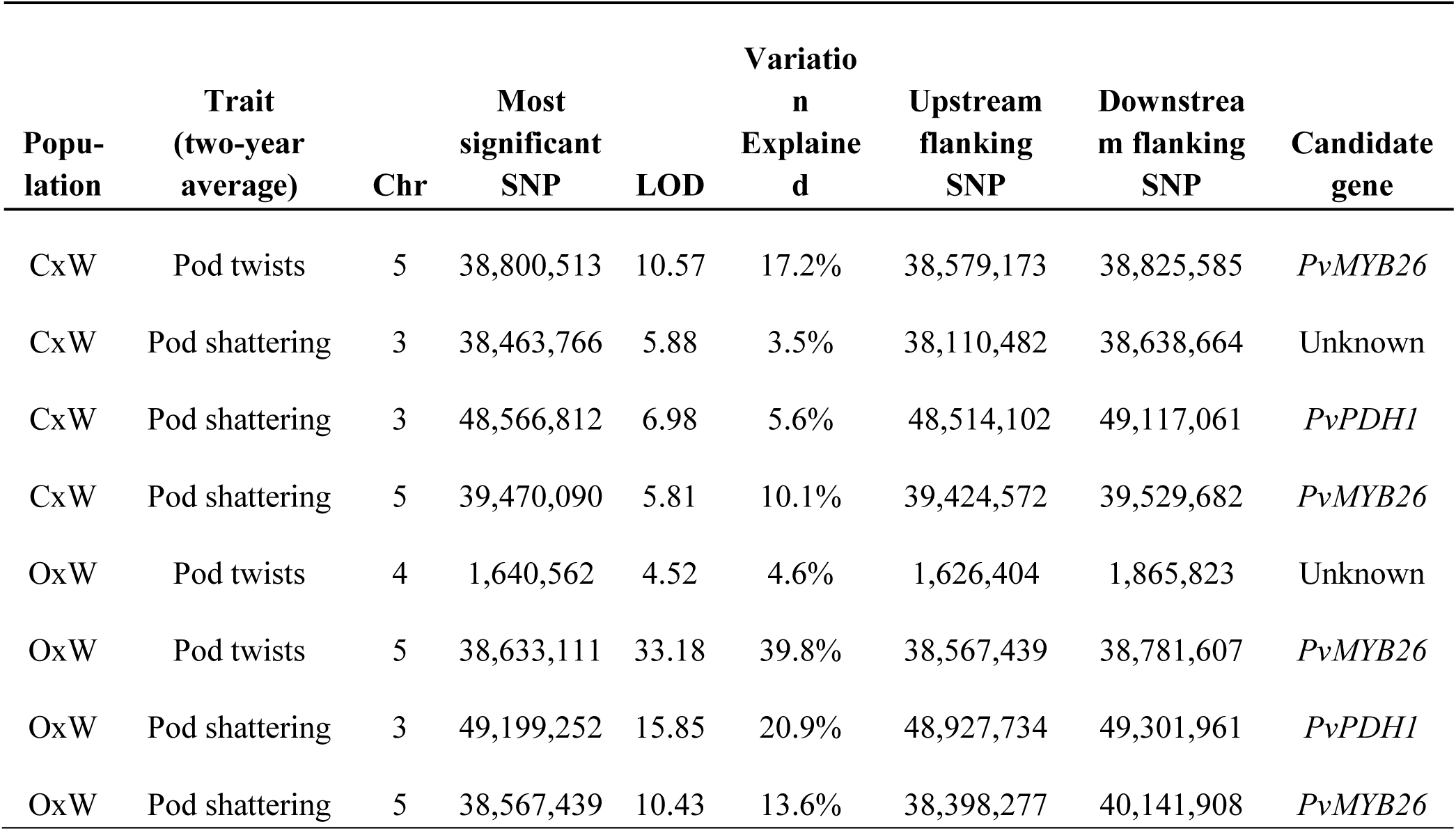
Significant QTL for pod shattering and pod twists in two wild x domesticated recombinant inbred populations.

**Supplementary Table 2.** *PvMYB26* allele frequencies in Wild, Middle American domesticated, and Andean domesticated common beans.

|  | 8 kb<br>deletion<br>present | 8 kb<br>deletion<br>absent | Total<br>genotyped | Percent with<br>deletion |
| --- | --- | --- | --- | --- |
| Wild | 2 | 88 | 90 | 2.2% |
| Middle American Diversity Panel | 109 | 1 | 110 | 99.1% |
| Andean Diversity Panel | 14 | 86 | 100 | 14% |

**Supplementary Table 3.** Middle American selective sweep statistics in a 600 kb region centered on *PvMYB26*. A 125 kb hard sweep was identified from 43,976,690 to 44,101,690 on Pv05 (5_593 genome v1.1). Data reflects a 600 kb region centered on the *PvMYB26*.

| Test statistic | 43,976,690-44,101,690 mean ±<br>SEM | 43,709,324-43,976,690;<br>44,101,690-44,309,325 mean<br>± SEM |
| --- | --- | --- |
| Nucleotide diversity (wild) | 0.0110 ± 0.000441 | 0.0109 ± 0.000245 |
| Nucleotide diversity (domesticated) | 0.00129 ± 0.000106 | 0.00570 ± 0.000194 |
| Nucleotide diversity–<br>(domesticated/wild) | 0.116 ± 0.00730 | 0.534 ± 0.0166 |
| Tajima’s D (wild) | 0.405 ± 0.0906 | 0.358 ± 0.0285 |
| Tajima’s D (domesticated) | -2.05 ± 0.0267 | 1.16 ± 0.0996 |
| $F_{ST}$ (wild vs domesticated) | 0.495 ± 0.0108 | 0.251 ± 0.00648 |

**Supplementary Table 4.** Andean selective sweep statistics selective sweep statistics in a 600 kb region centered on *PvMYB26*. No hard sweep was identifiable throughout the region. The coordinates 43,976,690 to 44,101,690 on Pv05 (5_593 genome v1.1) are the region of the hard sweep in Middle American lines for comparison.

| Test statistic | 43,976,690-44,101,690<br>mean ± SEM | 43,709,324-43,976,690;<br>44,101,690-44,309,325 mean ±<br>SEM |
| --- | --- | --- |
| Nucleotide diversity (wild) | 0.00302 ± 0.000268 | 0.00260 ± 0.000142 |
| Nucleotide diversity (domesticated) | 0.00240 ± 0.000111 | 0.00266 ± 0.000120 |
| Nucleotide diversity<br>(domesticated/wild) | 0.942 ± 0.0823 | 1.10 ± 0.0401 |
| Tajima’s D (wild) | -0.819 ± 0.0898 | -0.859 ± 0.0596 |
| Tajima’s D (domesticated) | -1.14 ± 0.0542 | -0.690 ± 0.0785 |
| *F<sub>ST</sub>* (wild vs domesticated) | 0.0900 ± 0.0177 | 0.118 ± 0.0125 |

**Supplementary Table 5.**
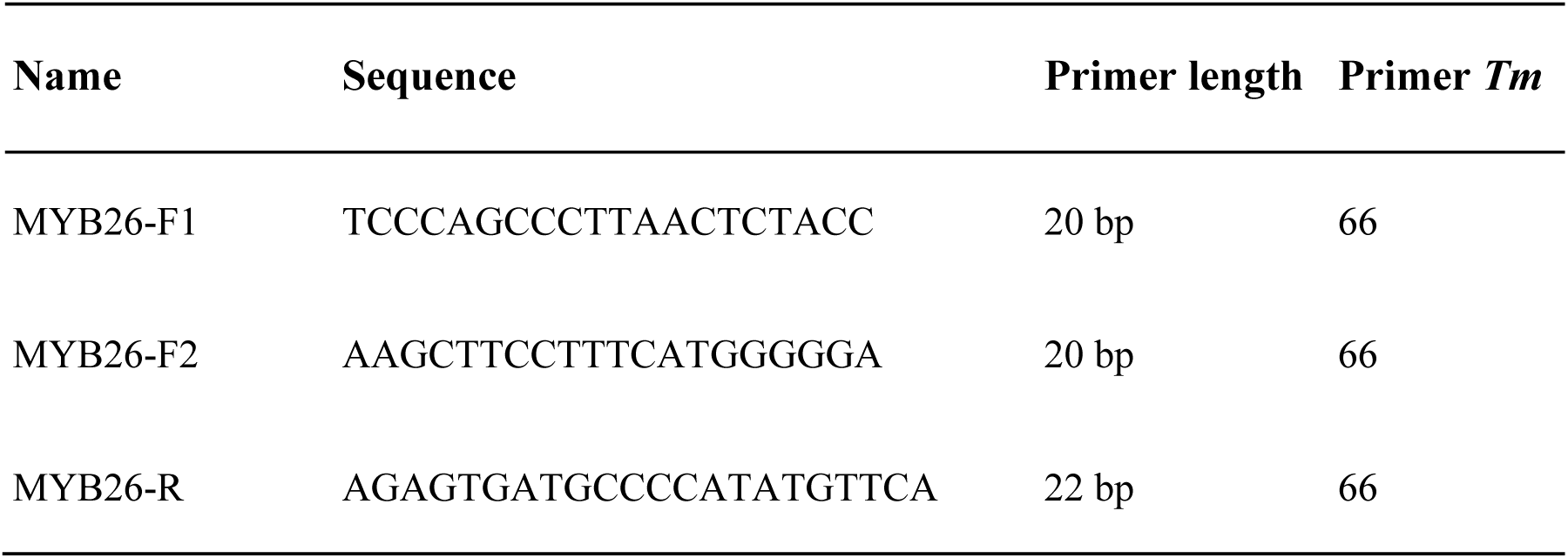
Co-dominant primers showing 8 kb deletion status in *PvMYB26*.

**Supplementary Table 6.**
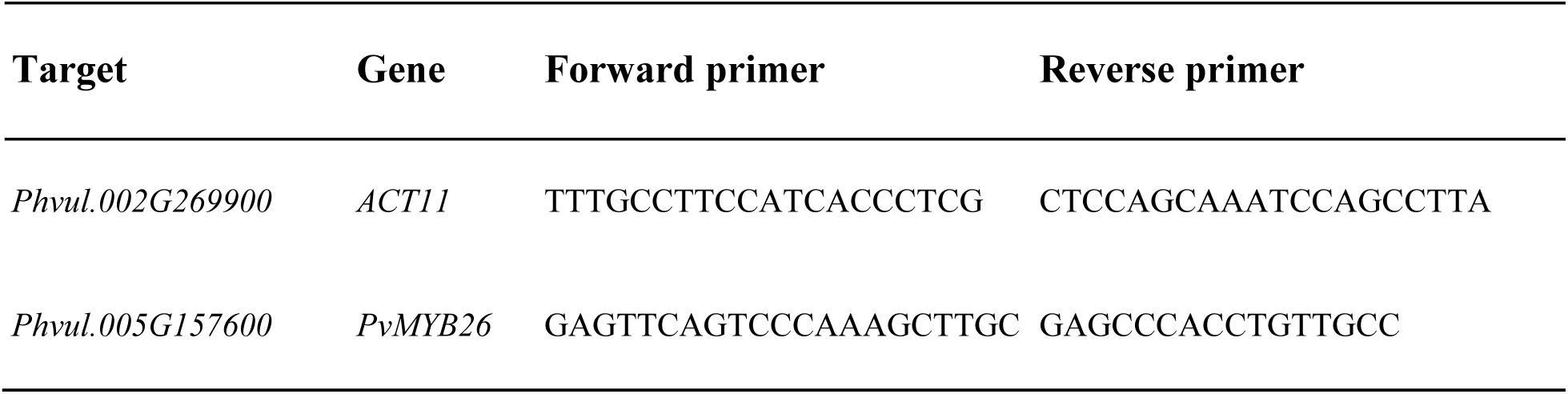
Primers for candidate gene (*PvMYB26*) and control gene (*Act11*)^100^ used for RT-qPCR.

## Supplementary Figures

**Supplementary Fig. 1.**
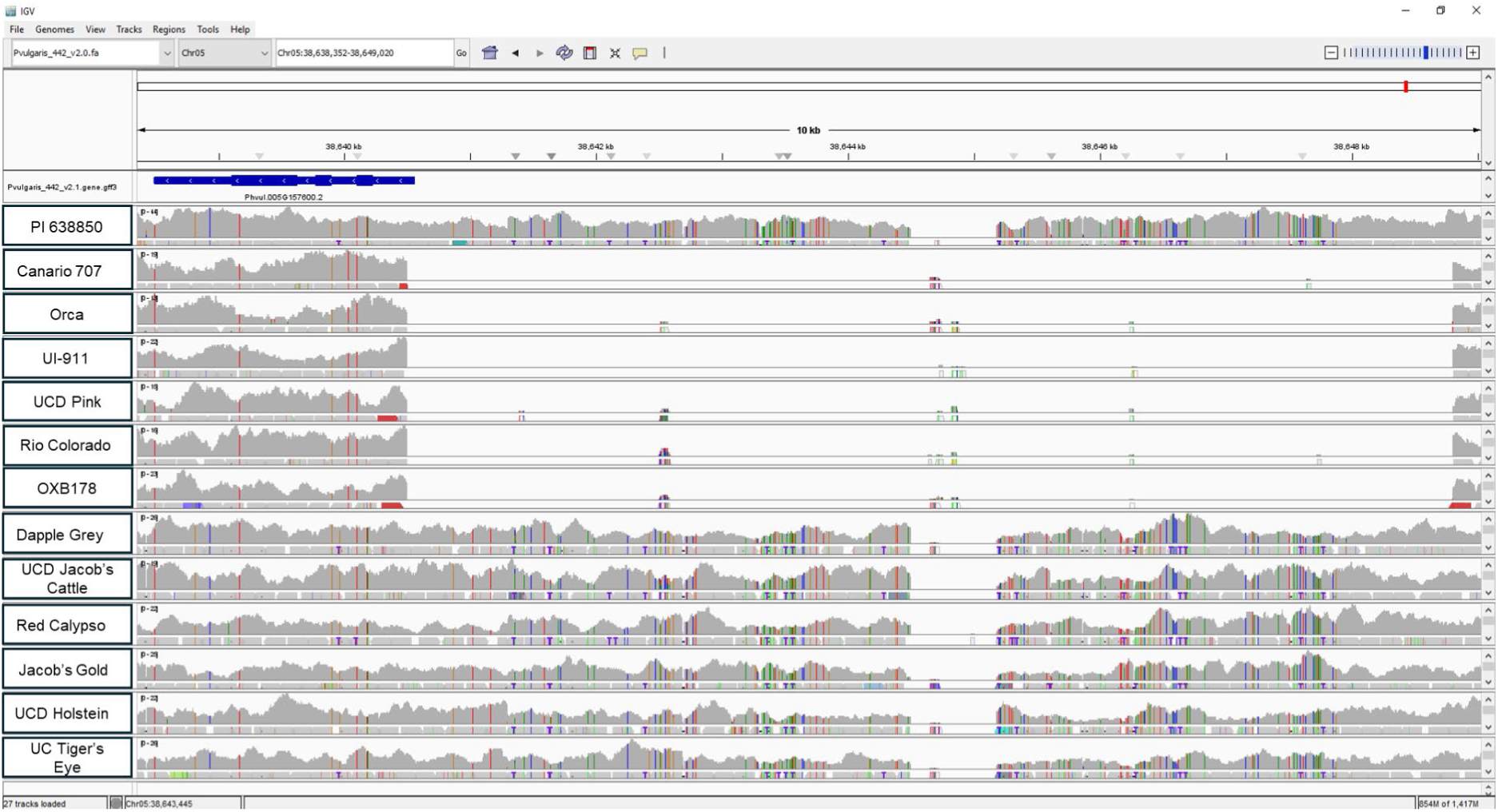
An IGV screenshot shows *PvMYB26* (*Phvul.005G157600*, transcribed region in blue at top left, exons thickened) and whole-genome-sequencing alignments with selected lines. The top track (PI 638850) is the wild parent of RIL populations and lacks the 8 kb deletion. The following six tracks show types with Middle American domesticated haplotypes, including the 8 kb deletion that eliminates the *PvMYB26* transcription start site and promoter. This includes the UC Canario 707 and Orca domesticated population parents. The last six tracks are types from the independently domesticated Andean gene pool, which lack the 8 kb deletion. Gray indicates the read depth in each accession at each base pair. A total absence of reads aligning to the reference indicates a deletion of the region in the sequenced experimental line. All sequences were aligned to the G19833 v2.1 genome^19^ (https://phytozome-next.jgi.doe.gov/info/Pvulgaris_v2_1).

**Supplementary Fig. 2.**
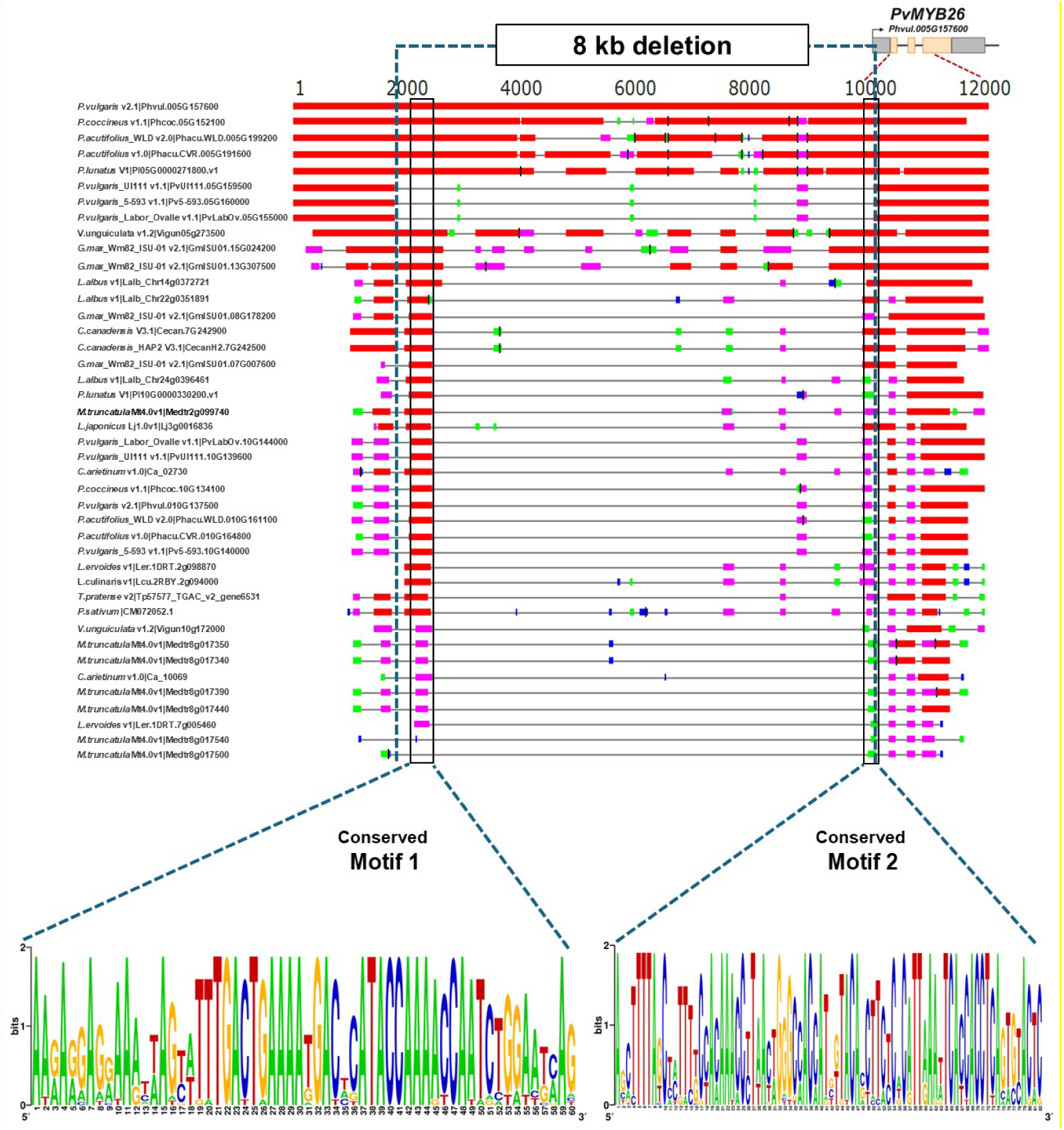
Two sequence motifs upstream of *PvMYB26* are highly conserved across the legume family, but are deleted in the domesticated RIL population parents and other domesticates with a Middle American haplotype at the gene. Motif 1 includes recognition sites of WRKY binding proteins (TTGAC)^101^ and MYC (bHLH) and MYB transcription factors (CANNTG and WAACCA)^102^. In Arabidopsis, the WRKY/MYC genes and auxin response factors (ARFs; ARF6/8) act upstream of the jasmonic acid (JA) biosynthesis and *MYB26* gene^103,104^. Disruption of this pathway results in suppression of *MYB26* expression and absence of secondary thickening, resulting in the production of non-opening anthers^103,104^. Motif 2 includes MYB recognition sites (TAACTG and YAACKG)^102^. Motif 2 includes an 82 bp sequence conserved in identical form across all *Phaseolus* species with available data (*P. acutifolius*, *P. coccineus*, *P. lunatus*, and wild-type *P. vulgaris*), as well as *Vigna angularis* and *Vigna unguiculata*, but 51 bp of which is deleted in Middle American domesticates.

**Supplementary Fig. 3.**
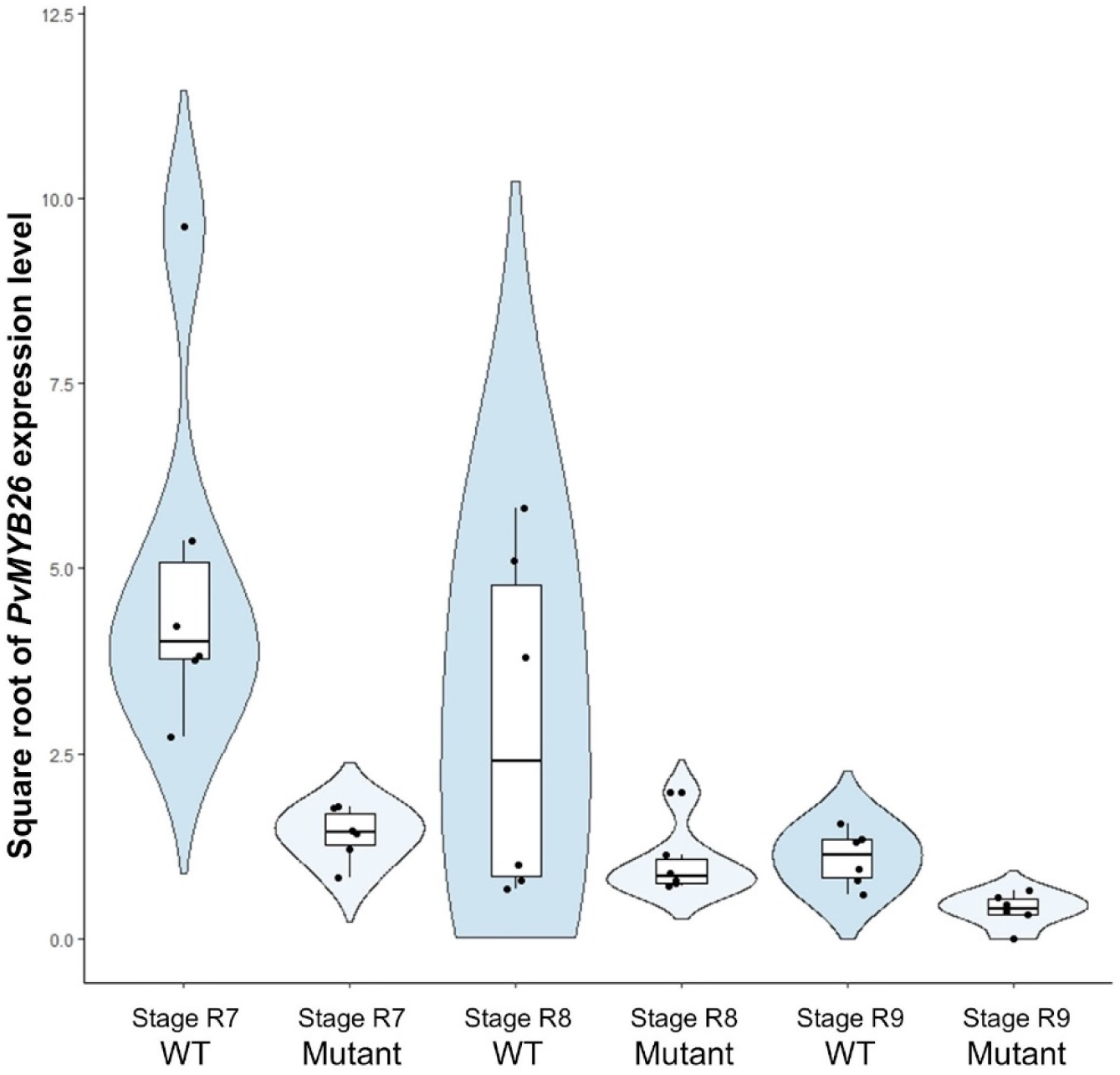
RNA-seq of wild and mutant genotypes^32^ across three pod maturity stages (Stage R7-9) showed significant differences in expression between *PvMYB26* mutants (accessions ICA Bunsi and SXB405) and non-mutants (accessions Midas and G12873) (*P* = 5.90 x 10^-5^). Expression was lower in mutants at every sampling time point.

**Supplementary Fig. 4.**
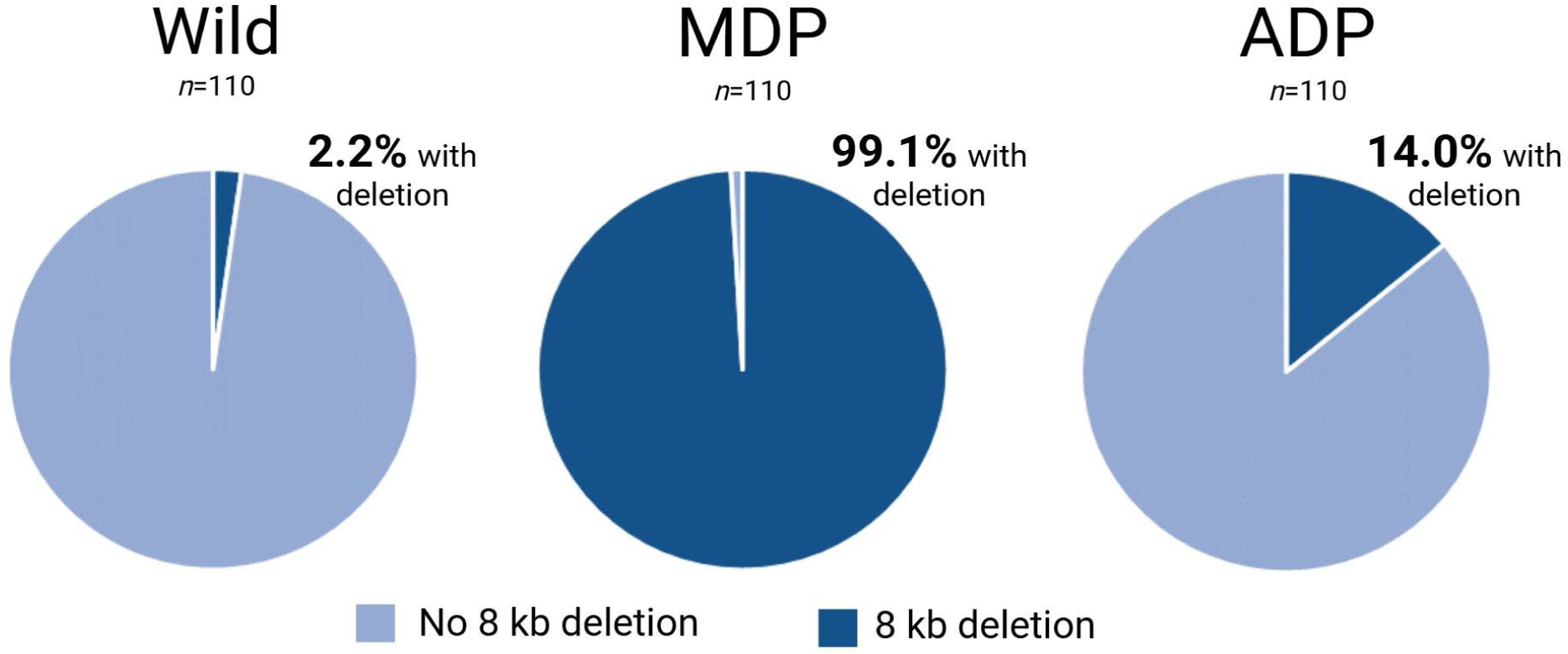
*PvMYB26* allele frequencies among wild common beans, members of the Middle American Diversity Panel (MDP), and the independently domesticated Andean Diversity Panel (ADP). The 8 kb *pvmyb26* deletion is nearly diagnostic for domestication status between wild and domesticated Middle American accessions of the species. While it originated in the Middle American gene pool, it has also been transferred to 14% of Andean domesticates.

**Supplementary Fig. 5.**
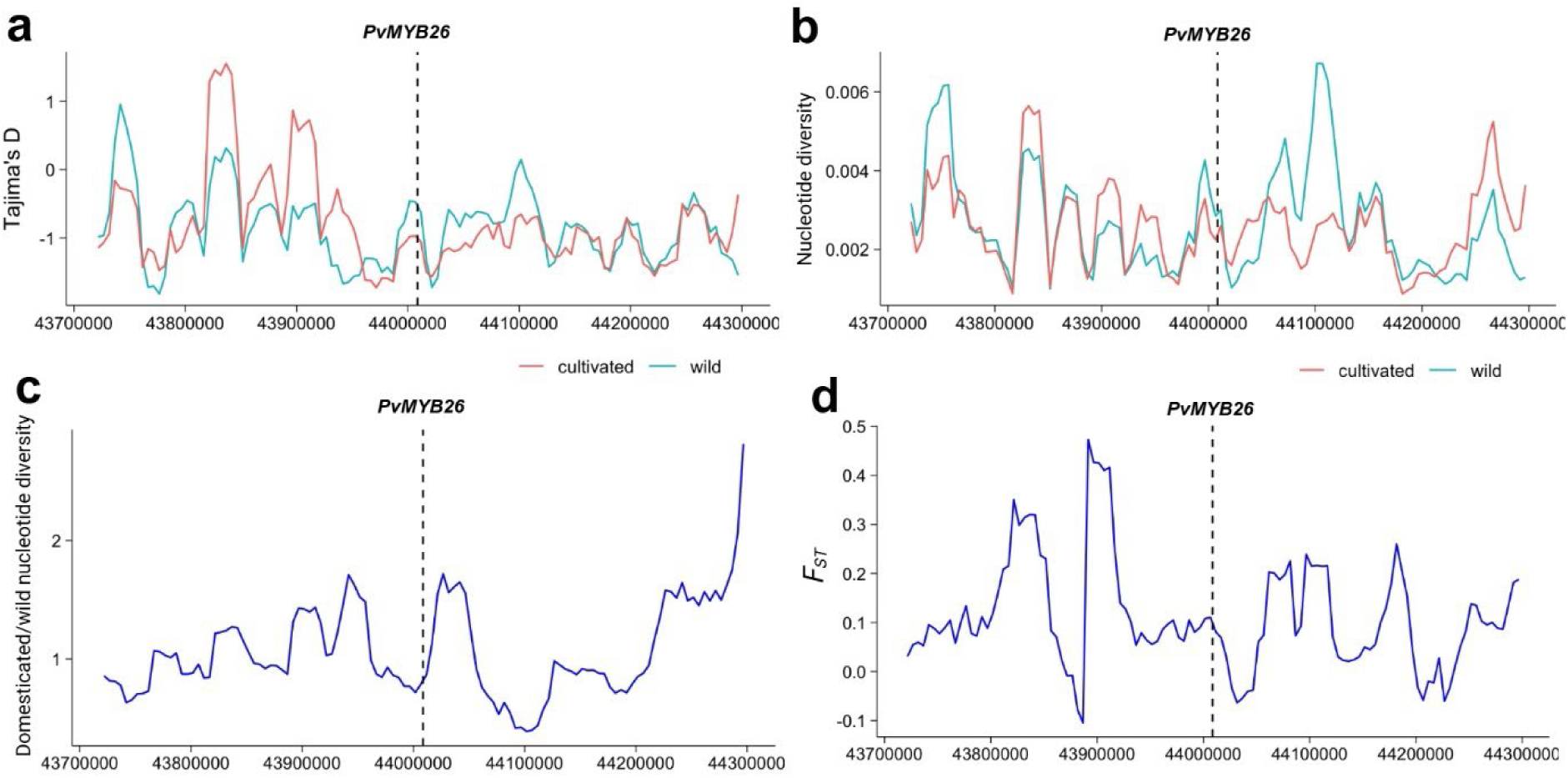
Population genetics statistics around *PvMYB26* show no selective sweep in Andean materials, in contrast to the hard sweep identified in Middle American materials. Calculated statistics include. **a,** Tajima’s D, **b-c,** nucleotide diversity, and **d,** *F_ST_*. Coordinates based on genome 5-593 v1.1.

